# Integrative Analysis of Proteomics and Metabolomics Reveals Impacts of *Sam2* Knockout on Autism Spectrum Disorders

**DOI:** 10.1101/2025.09.03.673986

**Authors:** Jaeho Ji, Minsook Ye, Cheol Soo Choi, Cheol-Hee Kim, Ahrum Son, Hyunsoo Kim

## Abstract

Neurokine Sam2 deficiency is associated with behavioral abnormalities including heightened anxiety and fear responses across evolutionarily conserved model organisms. Here, we employed an integrated proteomics and metabolomics approach using liquid chromatography-mass spectrometry (LC-MS) to elucidate molecular signatures associated with autism spectrum disorder (ASD) in *Sam2* knockout mice. Comparative analysis of blood plasma samples from *Sam2* knockout and wild-type mice revealed substantial alterations in both proteomic and metabolomic profiles. Proteomic analysis identified 68 differentially expressed proteins (comprising 102 peptides), with notable upregulation of complement component C1qc and downregulation of apolipoprotein A1 (Apoa1), implicating dysregulation of complement cascade pathways. Metabolomic profiling uncovered 15 significantly altered metabolites: nine upregulated species including D-glucuronic acid and 5,10-Methylenetetrahydrofolate, and six downregulated metabolites including folinic acid and acetate. Integrative pathway analysis revealed perturbations in glycolysis/gluconeogenesis and glycerophospholipid metabolism, providing mechanistic insights into the molecular consequences of Sam2 deficiency. These findings identify potential biomarkers for anxiety-related disorders and ASD while advancing our understanding of the complex interplay between genetic alterations and proteomic-metabolomic networks in neurodevelopmental conditions. Our results establish a foundation for developing targeted therapeutic interventions and highlight the utility of multi-omics approaches in dissecting the molecular basis of behavioral disorders.

## Introduction

Autism Spectrum Disorder (ASD) represents a heterogenous group of neurodevelopmental conditions arising from complex interactions between genetic and environmental factors, with a global prevalence of approximately 1 in 160 children.^1, 2^ While current therapeutic approaches rely predominantly on behavioral interventions and comprehensive healthcare management, the molecular mechanisms underlying ASD pathogenesis remain incompletely understood.

Recent investigations employing zebrafish and murine models have revealed that neurokine Sam2 deletion profoundly affect fear and anxiety responses, indicating an evolutionarily conserved role in behavioral regulation.^3, 4^ Notably, Sam2 dysfunction has been associated with intellectual disability and ASD-like phenotypes in zebrafish, prompting investigation into whether similar mechanisms operate in mammalian systems.

Additionally, SAM serves as a crucial cofactor in neurotransmitter biosynthesis, directly impacting mood regulation and cognitive function–domains frequently disrupted in ASD.^5-8^ Therefore, Sam2 deficiency could precipitate widespread cellular dysfunction contributing to ASD pathophysiology.

To elucidate the molecular consequences of Sam2 disruption, we employed an integrated proteomics and metabolomics approach to characterize the systemic alterations in Sam2 knockout mice. We hypothesized that Sam2 deletion would generate distinct molecular signatures in plasma, reflecting the biochemical perturbations underlying the observed behavioral phenotypes. By systematically comparing protein and metabolite profiles between Sam2 knockout and wild-type mice, we aimed to: (1) identify differentially expressed proteins and metabolites that could serve as biomarkers for early ASD diagnosis, (2) delineate the metabolic pathways disrupted by Sam2 deficiency, and (3) reveal potential therapeutic targets for intervention.

This study represents a critical step toward bridging the gap between behavioral manifestations and molecular mechanisms in Sam2-associated neurodevelopmental disorders. By integrating multi-omics approaches, we provide a comprehensive framework for understanding how Sam2 dysfunction contributes to ASD pathogenesis, ultimately advancing precision medicine approaches for affected individuals.

## Materials and Methods

### Reagents and Materials

Iodoacetamide was purchased from Sigma-Aldrich (St. Louis, MO, USA), Sequencing-grade modified trypsin was obtained from Promega (Madison, WI, USA). HPLC-grade water, formic acid, and acetonitrile were acquired from Thermo Fisher Scientific (Bremen, Germany). The Pierce^TM^ TOP14 Abundant Protein Depletion Spin Column was purchased from Thermo Fisher Scientific.

### Animals and Sample Collection

Blood samples were collected from Sam2 knockout (n=3) and wild-type (n=3) mice following institutional animal care guidelines. Plasma was separated by centrifugation, immediately flash-frozen in liquid nitrogen, and stored at -80^°^C until analysis. All samples were transported on dry ice to maintain sample integrity.

### Proteomic Analysis

#### Sample Preparation

Plasma proteins (100 μg) were quantified using the bicinchoninic acid (BCA) assay. High-abundance proteins were depleted using TOP14 columns according to the manufacturer’s protocol. Following depletion, protein concentrations were re-quantified, and samples were subjected to reduction with dithiothreitol (10 mM, 60^°^C, 30 min), alkylation with iodoacetamide (20 mM, room temperature, 30 min in darkness), and overnight digestion with trypsin (1:50 w/w enzyme:protein ratio) at 37^°^C. Peptides were desalted using C18 spin columns, concentrated by vacuum centrifugation, and stored at -80°C.

#### LC-MS/MS Analysis

Peptide analysis was performed using an Orbitrap Fusion mass spectrometer (Thermo Fisher Scientific) coupled with an UltiMate3000 RSLCnano system (Dionex). Peptides were separated on a nano HPLC capillary column (150 mm × 75 μm i.d., 3 μm particle size, Nikkyo Technos) using a 90-minute gradient of 5-35% acetonitrile in 0.1% formic acid at a flow rate of 300 nL/min.

### Metabolomic Analysis

#### Sample Preparation

Plasma samples (50 μL) were extracted with 500 μL cold methanol:water (1:1, v/v) by vortexing for 30 s. After centrifugation (15,000 × g, 10 min, 4^°^C), supernatants were collected and evaporated to dryness under vaccum. Dried extracts were reconstituted in 120 μL methanol:water (1:1, v/v) and centrifuged (10,000 × g, 10 min, 4^°^C) to remove particulates.

#### UPLC-Q-TOF MS Analysis

Metabolomic profiling was performed using an Agilent 1260 Infinity UPLC system coupled with an Agilent 6550 iFunnel Q-TOF mass spectrometer. Chromatographic separation was achieved on Hypersil GOLD C18 column (100×2.1 mm, 1.9 μm; Thermo Fisher Scientific) with a 3 μL injection volume. Data were acquired in both positive and negative electrospray ionization modes with a mass range of 50-1700 m/z.

### Data Processing and Statistical Analysis

#### Proteomic Data Analysis

Protein identification and label-free quantification (LFQ) were performed using MSFragger (v20.0) against the mouse UniProt database. LFQ intensities were quantile-normalized, and differential expression was assessed using Student’s t-test. Proteins were considered significantly altered with fold change >1.3 (upregulated) or <0.77 (downregulated) and false discovery rate (FDR)-adjusted p-value <0.05.

#### Metabolomic Data Analysis

Metabolite data were processed using MetaboAnalyst 6.0. Following quantile normalization, differential metabolites were identified based on: (1) variable importance in projection (VIP) scores >1 from OPLS-DA models, and (2) p-value <0.05 using Mann-Whitney U test.

#### Bioinformatics and Pathway Analysis

Principal component analysis (PCA), hierarchical clustering, and visualization (volcano plots, violin plots, heatmaps) were performed using R (v4.3.2). Pathway enrichment analysis was conducted using PROTEOMAPS (https://www.proteomaps.net/) for proteins and MetaboAnalyst for metabolites. Gene Ontology enrichment was performed using Metascape. Integration of proteomic and metabolomic data was achieved using MetScape (v3.1.3) in Cytoscape (v3.10.1). Statistical plots were generated using GraphPad Prism (v10.0).

## Results

### Selection of Protein Targets

The comprehensive workflow of our study is illustrated in **Fig. 1**. Plasma samples underwent sequential preparation for LC-MS analysis of both proteins and metabolites followed by data acquisition, processing, and statistical analysis using principal component analysis (PCA), orthogonal partial least squares discriminant analysis (OPLS-DA), and heatmap visualization. Subsequently, volcano plots and heatmaps were generated to identify differential expression patterns between *Sam2* KO and wild-type (WT) samples, culminating in an integrative multi-omics analysis of proteins and metabolites. In label-free quantification (LFQ) proteomics, raw intensity values reflect the relative abundance of peptides and proteins detected by mass spectrometry. To achieve normal distribution and enhance data interpretability, these raw intensity values were log_2_-transformed. The comparative analysis was restricted to peptide groups detected in at least 2 out of 3 biological replicates, thereby focusing on high-confidence peptides while excluding low-quality or irrelevant peptide groups **(Supplementary Fig. 1)**. To assess peptide overlap between experimental groups, a Venn diagram revealed 700 peptides in the KO group and 727 peptides in the WT group across all biological replicates **(Supplementary Fig. 2)**. Pearson correlation analysis of the complete dataset comprising 3,380 peptides, demonstrated distinct clustering patterns and significant differences in correlation coefficients between KO and WT groups **(Supplementary Figs. 3A & 3C)**. Following statistical filtering using Student’s t-test (*P* < 0.05), we identified 81 differentially expressed proteins represented by 131 significant peptides. Pearson correlation analysis of these filtered peptides maintained clear separation and significant differences between the experimental groups **(Supplementary Figs. 3B & 3D)**. To validate group discrimination, PCA was performed on the 131 statistically significant peptides (*P*-value < 0.05). The analysis revealed complete separation of WT and *Sam2* KO samples within their respective 95% confidence intervals, confirming significant biological differences between the two experimental groups **(Supplementary Fig. 4)**.

**Fig. 1.**
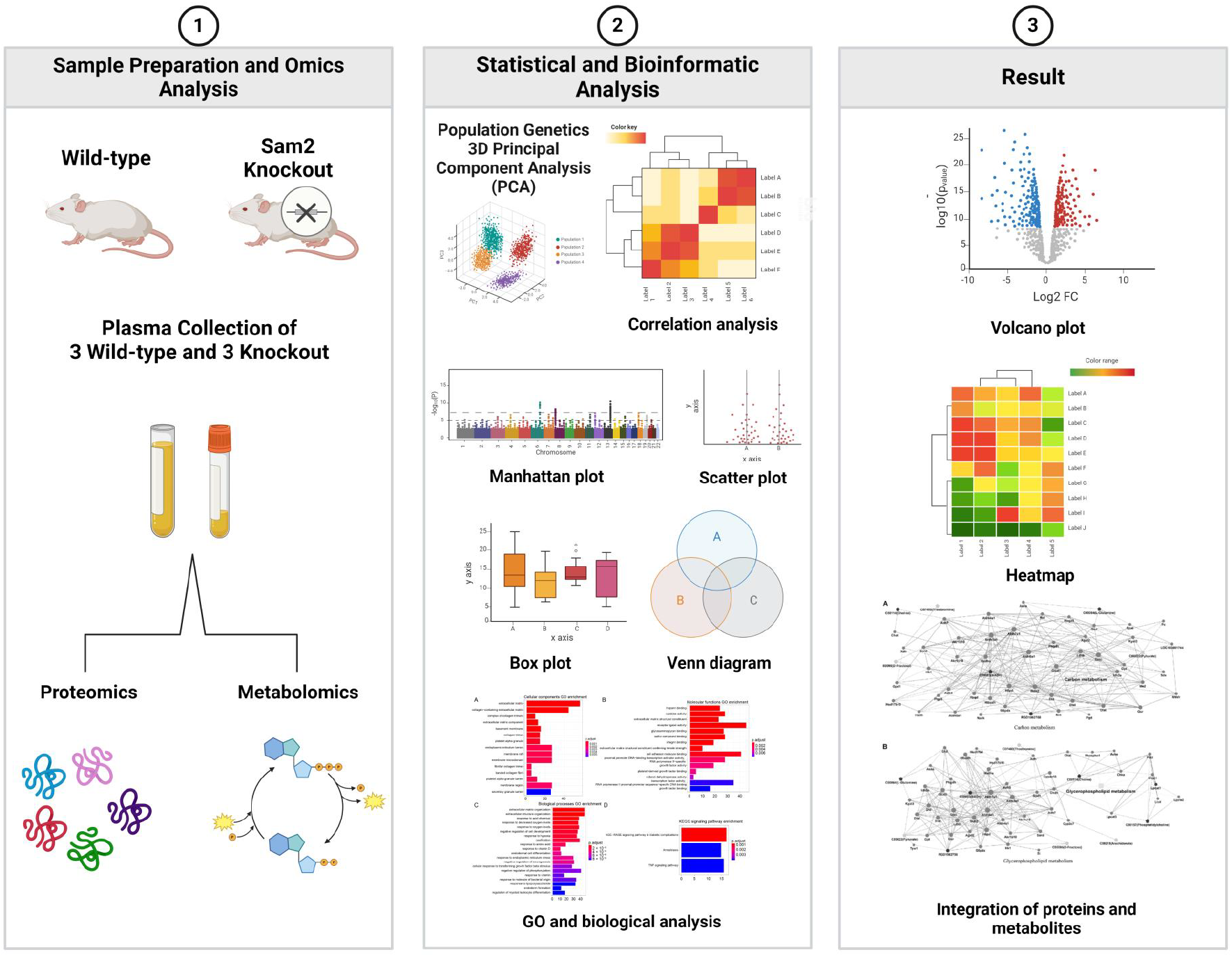
Overview of the Experimental Workflow. Schematic representation of the comparative multi-omics analysis workflow between *Sam2* knockout (KO) and wild-type (WT) mouse plasma samples. The workflow comprises three sequential phases: (1) sample preparation and LC-MS analysis for separate proteomic and metabolomic profiling; (2) data processing and statistical analysis including principal component analysis (PCA), orthogonal partial least squares discriminant analysis (OPLS-DA), and hierarchical clustering; and (3) differential expression analysis through volcano plots and heatmaps, followed by integrated multi-omics pathway analysis.

### Proteomic Profile Shift Between WT and *Sam2* KO

To characterize the proteomic alterations resulting from *Sam2* knockout, we performed hierarchical clustering analysis (HCA) on 101 significantly differentially expressed peptides (*P* < 0.05). The analysis revealed clear segregation between KO and WT groups, forming two distinct clusters that underscore the profound impact of Sam2 deletion on the proteome **(Fig. 2A)**. Pathway enrichment analysis using Proteomaps (https://www.proteomaps.net/) revealed contrasting biological signatures between the two clusters. Cluster 1 **(Fig. 2B)** and Cluster 2 **(Fig. 2C)** both demonstrated upregulation of immune system components and complement/coagulation cascades, with C1QC emerging as a consistently upregulated protein across both clusters. Conversely, endocrine system components and PPAR signaling pathway elements were significantly downregulated, particularly apolipoproteins. Quantitative assessment using volcano plot analysis (fold change threshold: > 1.3 for upregulation, < 0.77 for downregulation) confirmed the extensive proteomic remodeling, with 38 peptides upregulated and 63 downregulated among the 101 differentially expressed peptides **(Supplementary Fig. 5, Table 1)**. These integrated analyses–combining HCA, pathway enrichment, and quantitative proteomics–reveal that Sam2 deletion triggers a coordinated proteomic shift characterized by enhanced immune and coagulation responses coupled with suppressed endocrine and metabolic signaling. These findings provide mechanistic insights into how Sam2 deficiency fundamentally alters cellular physiology, with particular implications for immune function, hemostasis, and metabolic homeostasis.

**Fig. 2.**
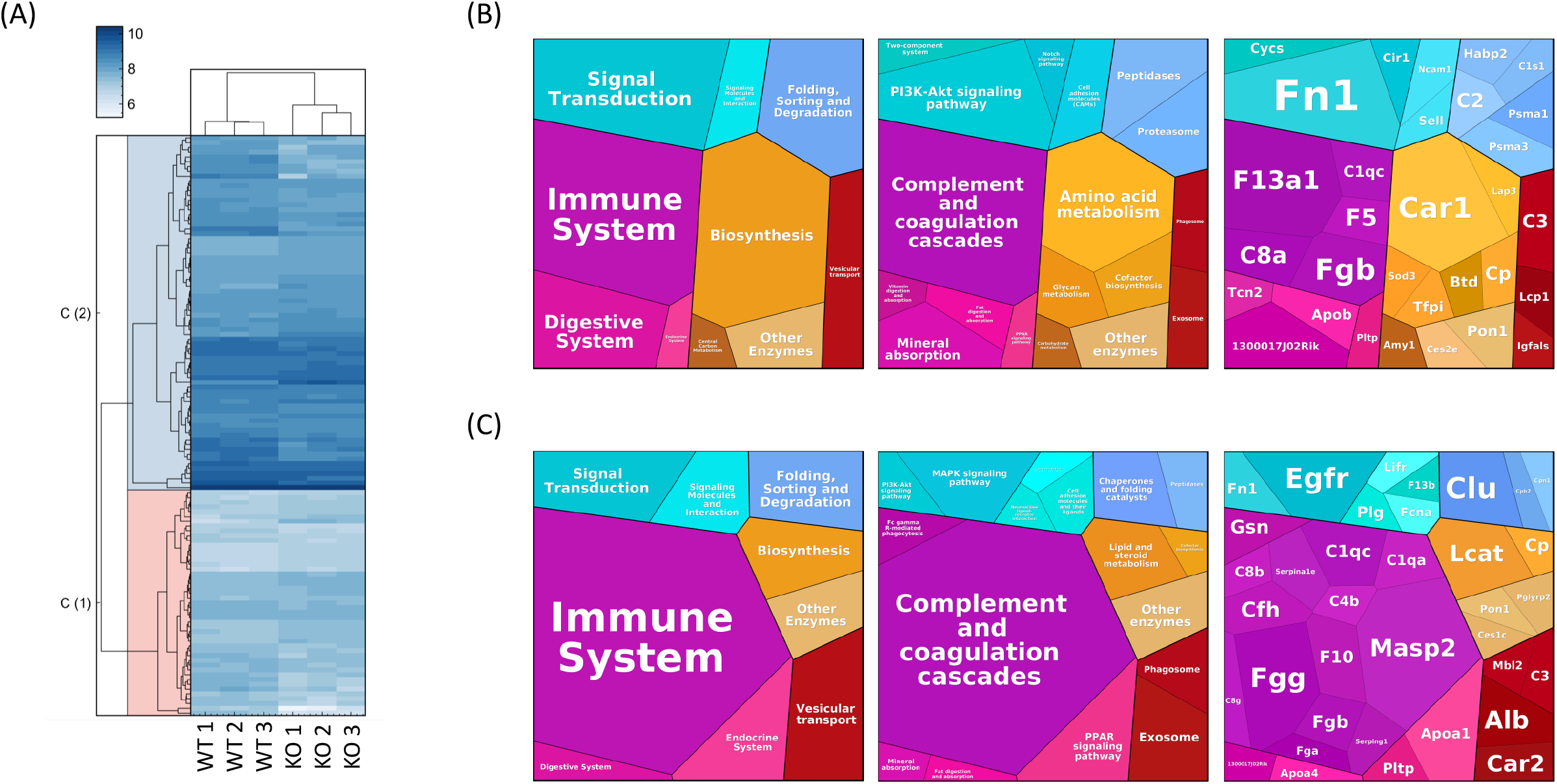
Hierarchical Clustering and KEGG Pathway Enrichment of Differentially Expressed Peptides. (A) Hierarchical cluster analysis of 131 significantly altered peptides (P < 0.05) demonstrating distinct clustering patterns between wild-type (WT) and knockout (KO) plasma samples. The heatmap visualization reveals two major clusters with contrasting expression profiles. (B, C) KEGG pathway enrichment analysis of Cluster 1 (C1) and Cluster 2 (C2) performed using Proteomaps, illustrating pathway-specific distributions of upregulated, downregulated, and total protein counts with statistical significance.

**Table 1.**
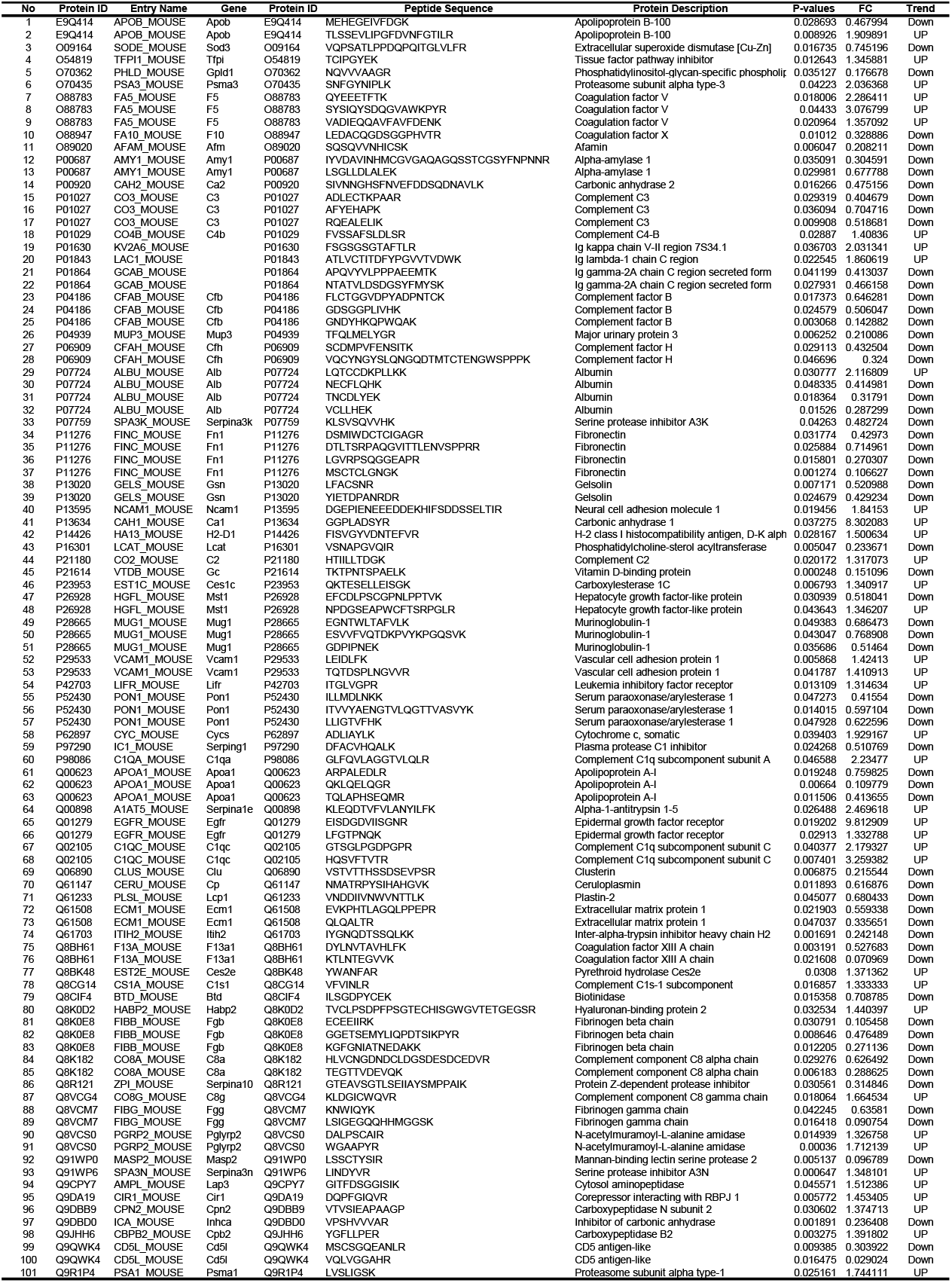
Comparison of Proteins Between Sam2 Gene KO and WT.

### Pathway Enrichment Analysis using Gene Ontology

The 101 identified peptides were subjected to GO enrichment analysis using Metascape^9^ with KEGG pathway annotations. Metascape generated a comprehensive protein-protein interaction (PPI) network, which was subsequently analyzed using the MCODE algorithm implemented in Cytoscape to identify densely connected regions representing potential protein complexes or functional modules **(Fig. 3A)**. The MCODE clustering analysis revealed distinct molecular complexes, with each cluster color-coded according to its functional identity. Notably, C1qc and Apoa1 were identified within two separate MCODE clusters, suggesting their involvement in distinct functional modules. Each MCODE cluster was annotated with enriched biological processes and pathways based on its constituent proteins. The enrichment results are presented in a Metascape bar graph displaying the top non-redundant enrichment clusters, with statistical significance represented by a discrete color scale **(Fig. 3B)**. These findings corroborated the Proteomaps results, showing significant enrichment in complement and coagulation cascades and complement activation biological processes.

**Fig. 3.**
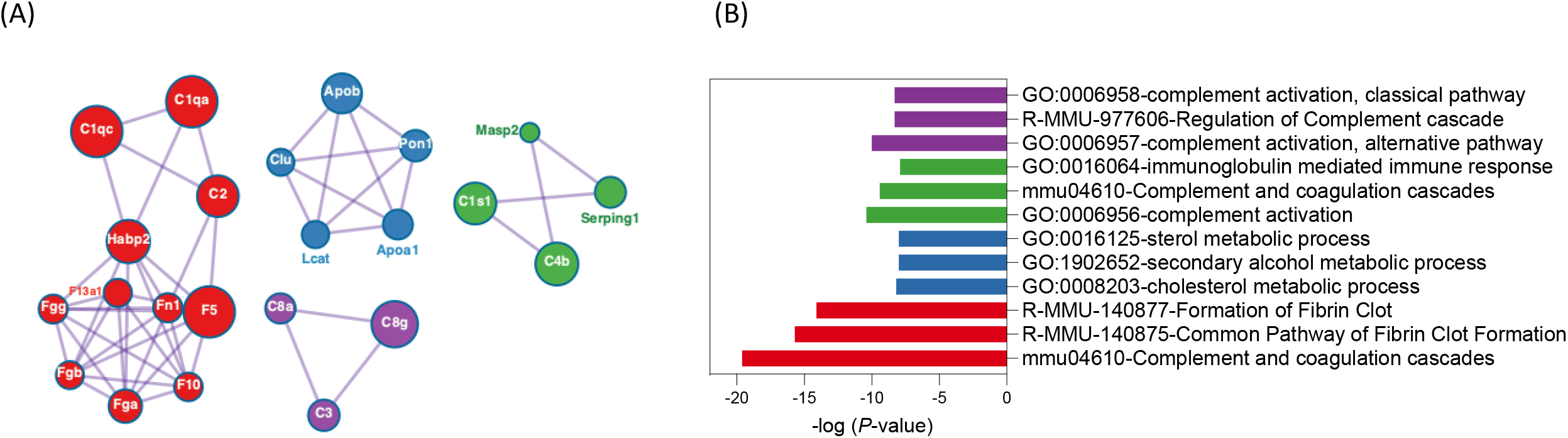
Gene Ontology Enrichment and Protein-Protein Interaction Network Analysis. (A) Metascape-generated Gene Ontology (GO) enrichment network displaying intra-cluster and inter-cluster relationships among enriched biological terms, with up to ten representative terms per cluster. The protein-protein interaction (PPI) network constructed from 121 candidate genes demonstrates MCODE-identified protein complexes color-coded by functional identity. C1qc and Apoa1 are highlighted within distinct cluster modules (red and blue, respectively), with node sizes reflecting the magnitude of differential expression between KO and WT groups. (B) Metascape bar graph presenting the top non-redundant enrichment clusters ranked by statistical significance, with discrete color scaling representing P-values.

To further validate these findings, we calculated the average LFQ intensities for C1qc and Apoa1 and generated bar plots to visualize differential expression between KO and WT groups (**Supplementary Fig. 6**). Statistical analysis revealed that C1qc was significantly upregulated in the KO group (*P* = 0.0237), while Apoa1 was significantly downregulated (*P* = 0.0025). In summary, PPI network analysis using Metascape with MCODE clustering successfully identified functionally distinct clusters containing C1qc and Apoa1. Pathway enrichment analysis highlighted the involvement of complement and coagulation cascades and complement activation processes. The differential expression patterns of these proteins–with C1qc upregulated and Apoa1 downregulated in KO versus WT conditions– suggest their opposing roles in the biological processes affected by the knockout phenotype.

### Identification of Differentially Expressed Metabolites

Metabolomic analysis detected a total of 7,933 features in positive ion mode and 3,445 features in negative ion mode. Following quality control filtering, 396 features were consistently identified across all biological replicates, from which 102 metabolites were successfully annotated in all plasma samples. Statistical analysis identified 15 metabolites with significant differential expression between groups (*P* < 0.05), which are presented in **Table 2** and visualized through heatmap analysis (**Supplementary Fig. 7**). The heatmap clustering pattern revealed distinct metabolic profiles between the experimental groups, with the KO group exhibiting elevated intensities for several metabolites compared to the WT group, indicating metabolic perturbations associated with the knockout phenotype.

**Table 2.** Comparison of Metabolites Between Sam2 Gene KO and WT.

| No | Metabolites | U stat value | P-value | VIP | Metabolic pathway | Adduct | pos/neg | m/z (mass to charge ratio) | Retention time (min) | Trend |
| --- | --- | --- | --- | --- | --- | --- | --- | --- | --- | --- |
| 1 | 5,10-Methylenetetrahydrofolate | 67 | 0.019 | 1.004 | One carbon pool by folate | M+2ACN+H | positive | 540.23465 | 643.7 | Upregulated |
| 2 | Folinic acid | 17 | 0.040 | 1.029 | One carbon pool by folate | M+ACN+Na | positive | 537.18517 | 473.0 | Downregulated |
| 3 | 10-Formyltetrahydrofolate | 17 | 0.040 | 1.029 | One carbon pool by folate | M+ACN+Na | positive | 537.18517 | 473.0 | Downregulated |
| 4 | Acetate | 9.5 | 0.007 | 1.480 | Glycolysis / Gluconeogenesis | 2M+H | positive | 121.04914 | 466.8 | Downregulated |
| 5 | beta-D-Glucuronoside | 8 | 0.005 | 1.619 | Pentose and glucuronate interconversions | M+Na | positive | 849.80747 | 49.8 | Downregulated |
| 6 | dGMP | 16.5 | 0.038 | 1.063 | Purine metabolism | M+H | positive | 348.07126 | 335.4 | Downregulated |
| 7 | Deoxyinosine | 66 | 0.027 | 1.063 | Purine metabolism | M+Na-2H | negative | 273.06025 | 866.9 | Upregulated |
| 8 | Pyridoxamine phosphate | 68 | 0.014 | 1.050 | Vitamin B6 metabolism | M+Na-2H | negative | 269.02878 | 45.6 | Upregulated |
| 9 | N-Acylsphingosine | 70.5 | 0.009 | 1.304 | Sphingolipid metabolism | M+H-2H<sub>2</sub>O | positive | 642.65094 | 447.7 | Upregulated |
| 10 | Sphingomyelin | 67.5 | 0.019 | 0.968 | Sphingolipid metabolism | M+ACN+2H | positive | 359.79982 | 877.8 | Upregulated |
| 11 | Glucosylceramide | 67.5 | 0.019 | 0.890 | Sphingolipid metabolism | M+2ACN+H | positive | 908.75770 | 564.8 | Upregulated |
| 12 | D-Glucuronic acid | 64.5 | 0.038 | 0.833 | Inositol phosphate metabolism | M+ACN+Na | positive | 258.05925 | 336.4 | Upregulated |
| 13 | D-Glucuronate | 64.5 | 0.038 | 0.833 | Ascorbate and aldarate metabolism | M+ACN+Na | positive | 258.05925 | 336.4 | Upregulated |
| 14 | 1-Acyl-sn-glycero-3-phosphocholine | 9 | 0.004 | 0.785 | Glycerophospholipid metabolism | M+2Na-H | positive | 560.27463 | 483.6 | Downregulated |
| 15 | CDP-choline | 69 | 0.011 | 0.847 | Glycerophospholipid metabolism | M+H-H<sub>2</sub>O | positive | 471.10453 | 816.7 | Upregulated |

### Metabolite Profile Shift Between WT and *Sam2* Gene KO

Prior to statistical analysis, all metabolite features were log_2_-transformed and normalized using z-transformation to ensure data comparability. Principle component analysis (PCA) demonstrated that plasma samples from both WT and *Sam2* KO groups clustered within the 95% confidence interval, confirming the reliability of the experimental design and data quality (**Fig. 4A**). Orthogonal partial least squares discriminant analysis (OPLS-DA) achieved complete separation between WT and KO groups, indicating robust metabolic differences between the two conditions (**Fig. 4B**). Model validation through permutation testing yielded high R^2^Y (79.3%) and Q^2^ (72.8%) values, demonstrating excellent model fit and predictive capability (**Supplementary Fig. 8A**). Further validation using 1,000 permutation tests confirmed the model’s statistical robustness, with enhanced R^2^Y (0.978) and Q^2^ (0.928) values, ruling out overfitting concerns (**Supplementary Fig. 8B**). Collectively, these validation metrics establish the OPLS-DA model as both statistically significant and analytically reliable. Variable importance in projection (VIP) analysis identified 15 metabolites with VIP scores ≥ 1.0, representing the most discriminatory features between groups (**Supplementary Fig. 8C**). Among these significantly altered metabolites, nine were upregulated in the KO group: D-glucuronic acid, 5,10-methylenetetrahydrofolate, D-glucuronate, deoxyinosine, CDP-choline, sphingomyelin, N-acylsphingosine, glucosylceramide, and pyridoxamine phosphate. Conversely, six metabolites were downregulated: folinic acid, 10-formyltetrahydrofolate, β-D-glucuronoside, dGMP, acetate, and 1-acyl-sn-glycero-3-phosphocholine (**Table 2**). This metabolic signature suggests substantial perturbations in folate metabolism, sphingolipid biosynthesis, and nucleotide metabolism pathways following *Sam2* gene knockout.

**Fig. 4.**
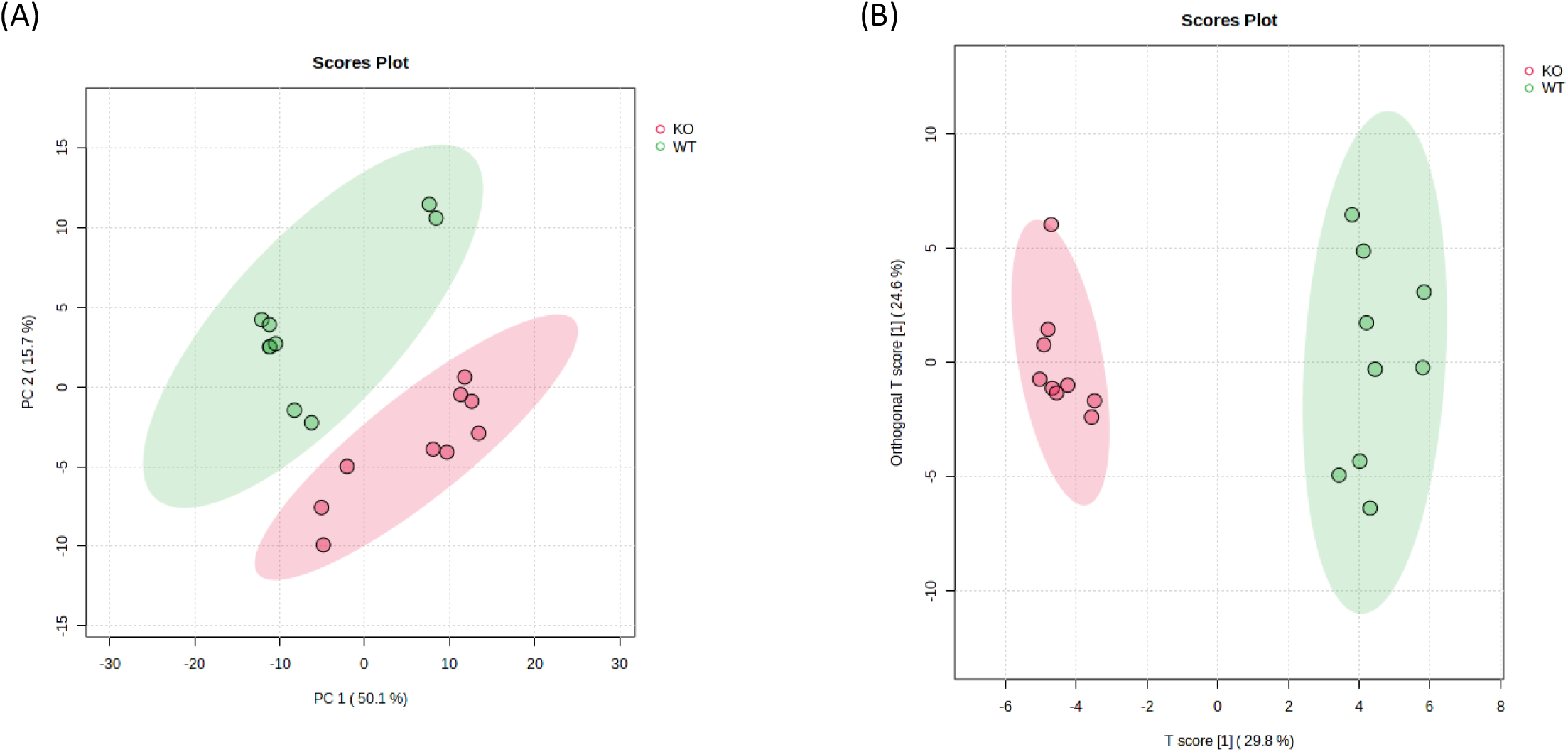
Multivariate Statistical Analysis of Metabolomic Profiles. Principal component analysis (PCA) and orthogonal partial least squares discriminant analysis (OPLS-DA) of 15 significantly altered metabolites demonstrate robust separation between experimental groups. (A) PCA score plot confirms data quality and reliability, with all plasma samples from both WT and *Sam2* KO groups clustering within the 95% confidence interval. (B) OPLS-DA analysis reveals complete discrimination between WT and KO metabolomic profiles, indicating substantial metabolic perturbations associated with *Sam2* gene knockout.

### Integration of Proteins and Metabolites

To elucidate the functional relationships between proteomic and metabolomic alterations, we performed integrated pathway analysis using the 101 significantly altered proteins and 15 differentially expressed metabolites (*P* < 0.05). Multi-omics integration analysis conducted through Metascape identified two significantly enriched pathways: glycolysis/gluconeogenesis and glycerophospholipid metabolism (**Fig. 5**). To validate these findings, we performed independent joint pathway analysis using Metaboanalyst, which corroborated the enrichment of the same metabolic pathways. The convergence of results from two distinct analytical platforms provides robust evidence for the involvement of glucose metabolism and phospholipid biosynthesis in the *Sam2* KO phenotype, demonstrating the reliability and biological significance of our integrated omics approach.

**Fig. 5.**
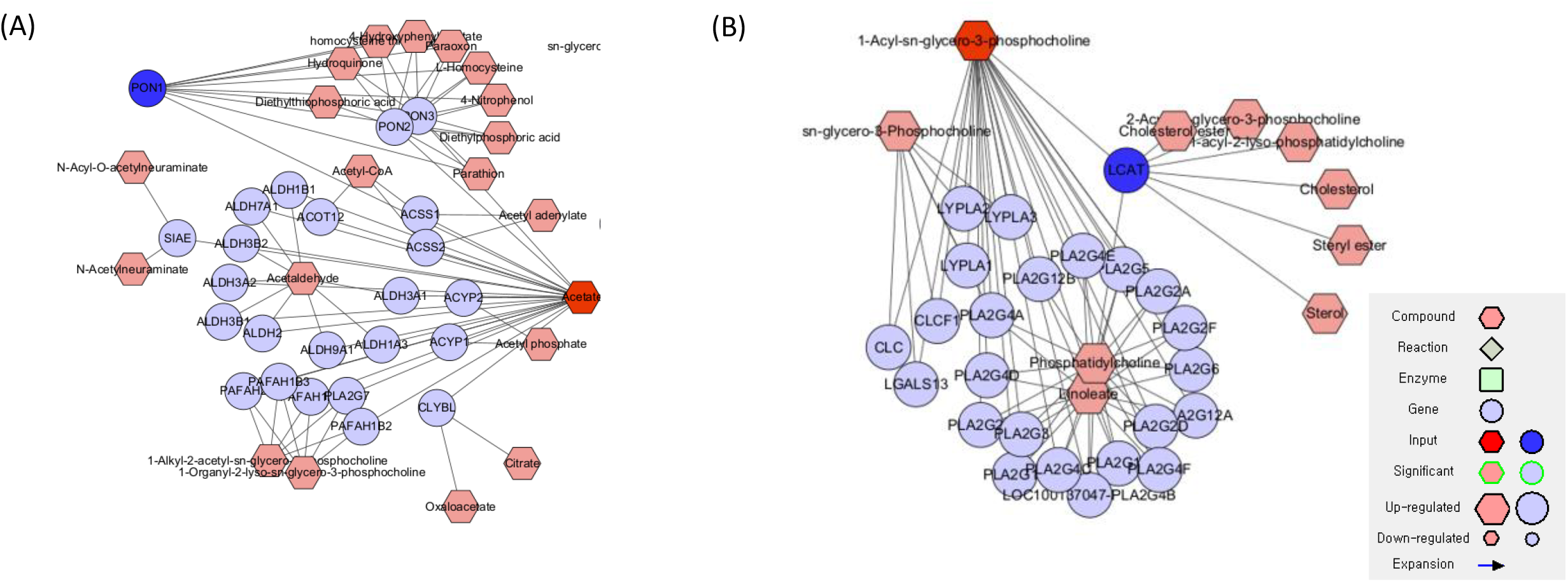
Integrated Multi-Omics Pathway Analysis. Metascape-mediated integration analysis of 101 differentially expressed proteins and 15 significantly altered metabolites (P < 0.05) identifies two major enriched metabolic pathways: glycolysis/gluconeogenesis and (B) glycerophospholipid metabolism. These pathways represent key metabolic networks significantly perturbed in the *Sam2* knockout phenotype, demonstrating the convergence of proteomic and metabolomic alterations in core cellular metabolism.

## Discussion

This study investigated the molecular consequences of *Sam2* gene knockout in a mouse model to elucidate potential mechanisms underlying ASD pathophysiology and identify candidate biomarkers for translational research. Our integrated proteomic and metabolomic analysis revealed distinctive molecular signatures that provide novel insights into ASD-related pathways. Proteomic analysis identified significant dysregulation of two key proteins: upregulation of C1qc and downregulation of Apoa1. The upregulation of C1qc is particularly noteworthy given the established role of C1q complement proteins in synaptic pruning during neurodevelopment.^10-14^ Aberrant synaptic pruning has been implicated in ASD pathophysiology, suggesting that C1qc could serve as a valuable biomarker for neurodevelopmental assessment in ASD. Conversely, the downregulation of Apoa1, a central regulator of lipid metabolism and transport, may contribute to the behavioral phenotypes observed in ASD, particularly heightened fear and anxiety responses.^15-18^ This finding aligns with emerging evidence linking lipid metabolic dysfunction to neuropsychiatric manifestations in autism.

Our metabolomic analysis revealed substantial alterations in metabolites associated with emotional regulation and neurotransmitter homeostasis.^19-24^ The upregulation of D-glucuronic acid, 5,10-methylenetetrahydrofolate, and CDP-choline, coupled with the downregulation of folinic acid and acetate, points to disrupted folate metabolism and gut-brain axis signaling.^25-30^ The reduction in acetate levels is particularly significant, as this short-chain fatty acid serves as a critical metabolic mediator produced by gut microbiota and has been directly linked to ASD-like behaviors.^31-33^ This finding suggests that *Sam2* knockout may indirectly influence gut microbiome-host metabolic interactions, contributing to the behavioral phenotype. The concurrent upregulation of deoxyinosine and pyridoxamine phosphate further supports dysregulation in purine metabolism and vitamin B6-dependent pathways, both of which are essential for neurotransmitter synthesis and oxidative stress management.^34-41^ Given vitamin B6’s crucial role in dopaminergic metabolism, these alterations may directly impact neurotransmitter balance and behavioral regulation in ASD.

A particularly novel finding was the identification of dysregulated ascorbate and aldarate metabolism, characterized by significant D-glucuronate upregulation. This pathway has received limited attention in ASD research, representing a previously unexplored metabolic dimension of the disorder. D-glucuronate plays essential roles in detoxification processes and vitamin C metabolism, potentially linking metabolic dysregulation to oxidative stress through a novel mechanistic pathway.^42, 43^ This discovery expands our understanding of ASD beyond traditionally studied metabolic networks and may reveal new therapeutic targets. The integration of proteomic and metabolomic datasets through multi-platform analysis consistently identified two core metabolic pathways: glycolysis/gluconeogenesis and glycerophospholipid metabolism. This convergence suggests that *Sam2* KO fundamentally alters cellular energy metabolism and membrane lipid homeostasis, processes that are critical for neuronal function and synaptic plasticity. The identification of these pathways provides a mechanistic framework for understanding how genetic perturbations translate into the complex behavioral phenotypes observed in ASD.

This comprehensive multi-omics analysis of *Sam2* KO mice has revealed a complex network of molecular alterations that collectively contribute to ASD-like phenotypes. The identification of C1qc as a potential biomarker, the discovery of novel metabolic pathway dysregulation, and the demonstration of gut-brain axis involvement provide multiple avenues for therapeutic intervention. Future studies should focus on validating these findings in human ASD populations and exploring the therapeutic potential of targeting the identified metabolic pathways. The integration of genetics, proteomics, and metabolomics demonstrated here establishes a robust framework for advancing precision medicine approaches in autism spectrum disorders.

## Acknowledgement

This work was supported by the National Research Foundation of Korea (NRF) grant funded by the Korean government (MSIT) (RS-2023-00209456) and the Korea Basic Science Institute (National Research Facilities and Equipment Center) grant funded by the Korean government (MSIT) (RS-2024-00402298). Additional support was provided by the Korea National Institute of Health (KNIH) research project (2024ER051900). We acknowledge the use of Claude Opus 4 solely for linguistic refinement and grammatical corrections in manuscript preparation. All scientific content, data analysis, and intellectual contributions presented herein were developed independently by the authors without the use of generative AI tools. Figure 1 was created with BioRender.com (accessed 1 July 2025).

## Supplementary Figure Legend

**Supplementary Fig. 1. Quality Assessment of Quantified Peptide Intensities**.

Violin plot comparing log-transformed peptide intensities between knockout (KO) and wild-type (WT) groups. The analysis includes peptides quantified in at least two out of three technical replicates, with a median log-transformed intensity value of 8. The plot demonstrates the distribution and quality of peptide quantification across experimental groups.

**Supplementary Fig. 2. Peptide Identification Overlap Analysis Across Biological Replicates**.

Venn diagrams illustrating peptide identification consistency across three biological replicates. (A) *Sam2* KO samples: approximately 2,000 peptides identified per replicate, with 700 peptide groups consistently detected across all three replicates and 1,434 peptide groups identified in at least two replicates. (B) Wild-type samples: 1,427 peptide groups consistently identified across all three biological replicates, demonstrating robust peptide detection.

**Supplementary Fig. 3. Correlation Analysis of Proteomic Data Quality and Statistical Filtering**.

Pearson correlation matrices demonstrating data reproducibility and statistical filtering effects. (A) Correlation analysis of LFQ intensities for 3,380 peptides quantified in at least two replicates, showing clear separation between WT and KO samples. (B) Correlation matrix of 131 statistically significant peptides (P < 0.05) after t-test filtering, confirming robust group discrimination. (C) Biological replicate correlation for the complete dataset of 359 proteins based on LFQ intensities. (D) Correlation analysis of 81 significantly altered proteins (P < 0.05) after statistical filtering, demonstrating maintained data quality post-filtering.

**Supplementary Fig. 4. Principal Component Analysis of Significantly Altered Peptides**.

PCA score plot of 131 significantly altered peptides (P < 0.05) demonstrating clear separation between KO (red) and WT (blue) experimental groups. The first two principal components account for 81.58% and 5.45% of the total variance, respectively, indicating robust multivariate discrimination between experimental conditions.

**Supplementary Fig. 5. Volcano Plot Analysis of Differential Peptide Expression**.

Volcano plot visualization of 101 significantly altered peptides (P < 0.05) between KO and WT groups. Red points represent significantly upregulated peptides (fold change > 1.3), while blue points indicate significantly downregulated peptides (fold change < 0.77) in the KO group relative to WT controls.

**Supplementary Fig. 6. Quantitative Validation of Key Cluster-Associated Proteins**.

Bar plots displaying LFQ intensities of representative proteins from distinct clusters. (A) C1qc expression levels from Cluster 1, showing significant upregulation in KO compared to WT samples (P < 0.05). (B) Apoa1 expression levels from Cluster 2, demonstrating significant downregulation in KO relative to WT samples (P < 0.05). Error bars represent standard error of the mean.

**Supplementary Fig. 7. Hierarchical Clustering of Significantly Altered Metabolites**.

Heatmap visualization of 15 significantly altered metabolites (P < 0.05, n = 15) between KO and WT groups. Color intensity represents normalized metabolite abundance, with red indicating upregulation and blue indicating downregulation in the KO group relative to WT controls. Hierarchical clustering groups metabolites with similar expression patterns, revealing predominant upregulation (darker red) in the KO group.

**Supplementary Fig. 8. Statistical Validation and Variable Selection for OPLS-DA Model**.

Comprehensive validation of the OPLS-DA model through multiple statistical approaches. (A) Permutation test validation demonstrating model robustness (R^2^Y = 79.3%, Q^2^ = 72.8%). (B) Enhanced validation through 1,000 permutation tests confirming model reliability (R^2^Y = 0.978, Q^2^ = 0.928, P = 0.001). (C) Variable importance in projection (VIP) plot identifying 15 metabolites with VIP scores ≥ 1.0, representing the most discriminatory features between experimental groups. Collectively, these analyses confirm the statistical significance and predictive reliability of the OPLS-DA model.

## References

1. van ‘t Hof M, Tisseur C, van Berckelear-Onnes I, van Nieuwenhuyzen A, Daniels AM, Deen M et al. Age at autism spectrum disorder diagnosis: A systematic review and meta-analysis from 2012 to 2019. Autism 2021; 25(4): 862–873.

2. Salari N, Rasoulpoor S, Rasoulpoor S, Shohaimi S, Jafarpour S, Abdoli N et al. The global prevalence of autism spectrum disorder: a comprehensive systematic review and meta-analysis. Ital J Pediatr 2022; 48(1): 112.

3. Ariyasiri K, Choi TI, Kim OH, Hong TI, Gerlai R, Kim CH. Pharmacological (ethanol) and mutation (sam2 KO) induced impairment of novelty preference in zebrafish quantified using a new three-chamber social choice task. Prog Neuropsychopharmacol Biol Psychiatry 2019; 88: 53–65.

4. Choi JH, Jeong YM, Kim S, Lee B, Ariyasiri K, Kim HT et al. Targeted knockout of a chemokine-like gene increases anxiety and fear responses. Proc Natl Acad Sci U S A 2018; 115(5): E1041–E1050.

5. Rea V, Van Raay TJ. Using Zebrafish to Model Autism Spectrum Disorder: A Comparison of ASD Risk Genes Between Zebrafish and Their Mammalian Counterparts. Front Mol Neurosci 2020; 13: 575575.

6. Henriquez Martinez A, Avila LC, Pulido MA, Ardila YA, Akle V, Bloch NI. Age-Dependent Effects of Chronic Stress on Zebrafish Behavior and Regeneration. Front Physiol 2022; 13: 856778.

7. Choi TY, Choi TI, Lee YR, Choe SK, Kim CH. Zebrafish as an animal model for biomedical research. Exp Mol Med 2021; 53(3): 310–317.

8. Remines M, Schoonover MG, Knox Z, Kenwright K, Hoffert KM, Coric A et al. Profiling the compendium of changes in Saccharomyces cerevisiae due to mutations that alter availability of the main methyl donor S-Adenosylmethionine. G3 (Bethesda) 2024; 14(4).

9. Zhou Y, Zhou B, Pache L, Chang M, Khodabakhshi AH, Tanaseichuk O et al. Metascape provides a biologist-oriented resource for the analysis of systems-level datasets. Nat Commun 2019; 10(1): 1523.

10. Dias A, Santos D, Coelho T, Alves-Ferreira M, Sequeiros J, Alonso I et al. C1QA and C1QC modify age-at-onset in familial amyloid polyneuropathy patients. Ann Clin Transl Neurol 2019; 6(4): 748–754.

11. Carbutt S, Duff J, Yarnall A, Burn DJ, Hudson G. Variation in complement protein C1q is not a major contributor to cognitive impairment in Parkinson’s disease. Neurosci Lett 2015; 594: 66–69.

12. Magdalon J, Mansur F, Teles ESAL, de Goes VA, Reiner O, Sertie AL. Complement System in Brain Architecture and Neurodevelopmental Disorders. Front Neurosci 2020; 14: 23.

13. Zhang W, Chen Y, Pei H. C1q and central nervous system disorders. Front Immunol 2023; 14: 1145649.

14. Tran MTN, Hamada M, Jeon H, Shiraishi R, Asano K, Hattori M et al. MafB is a critical regulator of complement component C1q. Nat Commun 2017; 8(1): 1700.

15. Li Q, Shi Y, Li X, Yang Y, Zhang X, Xu L et al. Proteomic-Based Approach Reveals the Involvement of Apolipoprotein A-I in Related Phenotypes of Autism Spectrum Disorder in the BTBR Mouse Model. Int J Mol Sci 2022; 23(23).

16. Tierney E, Remaley AT, Thurm A, Jager LR, Wassif CA, Kratz LE et al. Sterol and lipid analyses identifies hypolipidemia and apolipoprotein disorders in autism associated with adaptive functioning deficits. Transl Psychiatry 2021; 11(1): 471.

17. Yao F, Zhang K, Feng C, Gao Y, Shen L, Liu X et al. Protein Biomarkers of Autism Spectrum Disorder Identified by Computational and Experimental Methods. Front Psychiatry 2021; 12: 554621.

18. Corbett BA, Kantor AB, Schulman H, Walker WL, Lit L, Ashwood P et al. A proteomic study of serum from children with autism showing differential expression of apolipoproteins and complement proteins. Mol Psychiatry 2007; 12(3): 292–306.

19. Osman A, Mervosh NL, Strat AN, Euston TJ, Zipursky G, Pollak RM et al. Acetate supplementation rescues social deficits and alters transcriptional regulation in prefrontal cortex of Shank3 deficient mice. Brain Behav Immun 2023; 114: 311–324.

20. Frye RE, Slattery J, Delhey L, Furgerson B, Strickland T, Tippett M et al. Folinic acid improves verbal communication in children with autism and language impairment: a randomized double-blind placebo-controlled trial. Mol Psychiatry 2018; 23(2): 247–256.

21. Renard E, Leheup B, Gueant-Rodriguez RM, Oussalah A, Quadros EV, Gueant JL. Folinic acid improves the score of Autism in the EFFET placebo-controlled randomized trial. Biochimie 2020; 173: 57–61.

22. Peralta-Marzal LN, Prince N, Bajic D, Roussin L, Naudon L, Rabot S et al. The Impact of Gut Microbiota-Derived Metabolites in Autism Spectrum Disorders. Int J Mol Sci 2021; 22(18).

23. Zhang A, Yan G, Zhou X, Wang Y, Han Y, Guan Y et al. High resolution metabolomics technology reveals widespread pathway changes of alcoholic liver disease. Mol Biosyst 2016; 12(1): 262–273.

24. Siracusano M, Arturi L, Riccioni A, Noto A, Mussap M, Mazzone L. Metabolomics: Perspectives on Clinical Employment in Autism Spectrum Disorder. Int J Mol Sci 2023; 24(17).

25. Naviaux JC, Schuchbauer MA, Li K, Wang L, Risbrough VB, Powell SB et al. Reversal of autism-like behaviors and metabolism in adult mice with single-dose antipurinergic therapy. Transl Psychiatry 2014; 4(6): e400.

26. Gevi F, Belardo A, Zolla L. A metabolomics approach to investigate urine levels of neurotransmitters and related metabolites in autistic children. Biochim Biophys Acta Mol Basis Dis 2020; 1866(10): 165859.

27. Dai S, Lin J, Hou Y, Luo X, Shen Y, Ou J. Purine signaling pathway dysfunction in autism spectrum disorders: Evidence from multiple omics data. Front Mol Neurosci 2023; 16: 1089871.

28. Geryk J, Krsicka D, Vlckova M, Havlovicova M, Macek M, Jr., Kremlikova Pourova R. The Key Role of Purine Metabolism in the Folate-Dependent Phenotype of Autism Spectrum Disorders: An In Silico Analysis. Metabolites 2020; 10(5).

29. Cappuccio G, Donti T, Pinelli M, Bernardo P, Bravaccio C, Elsea SH et al. Sphingolipid Metabolism Perturbations in Rett Syndrome. Metabolites 2019; 9(10).

30. Hussain G, Wang J, Rasul A, Anwar H, Imran A, Qasim M et al. Role of cholesterol and sphingolipids in brain development and neurological diseases. Lipids Health Dis 2019; 18(1): 26.

31. Mehta D, Davis M, Epstein AJ, Wensel B, Grinnell T, Thach A et al. Comparative economic outcomes in patients with focal seizures initiating eslicarbazepine acetate versus brivaracetam as their first adjunctive ASD. J Med Econ 2021; 24(1): 939–948.

32. Qin Y, Zhang XY, Liu Y, Ma Z, Tao S, Li Y et al. Downregulation of mGluR1-mediated signaling underlying autistic-like core symptoms in Shank1 P1812L-knock-in mice. Transl Psychiatry 2023; 13(1): 329.

33. Levi Mortera S, Vernocchi P, Basadonne I, Zandona A, Chierici M, Durighello M et al. A metaproteomic-based gut microbiota profiling in children affected by autism spectrum disorders. J Proteomics 2022; 251: 104407.

34. Huh E, Choi JG, Lee MY, Kim JH, Choi Y, Ju IG et al. Peripheral metabolic alterations associated with pathological manifestations of Parkinson’s disease in gut-brain axis-based mouse model. Front Mol Neurosci 2023; 16: 1201073.

35. Liu X, Liu M, Liu H, Yuan H, Wang Y, Chen X et al. Comprehensive brain tissue metabolomics and biological network technology to decipher the mechanism of hydrogen-rich water on Radiation-induced cognitive impairment in rats. BMC Mol Cell Biol 2023; 24(1): 30.

36. Liu X, Lin J, Zhang H, Khan NU, Zhang J, Tang X et al. Oxidative Stress in Autism Spectrum Disorder-Current Progress of Mechanisms and Biomarkers. Front Psychiatry 2022; 13: 813304.

37. Berridge MJ. The Inositol Trisphosphate/Calcium Signaling Pathway in Health and Disease. Physiol Rev 2016; 96(4): 1261–1296.

38. Peng L, Chen L, Wan J, Liu W, Lou S, Shen Z. Single-cell transcriptomic landscape of immunometabolism reveals intervention candidates of ascorbate and aldarate metabolism, fatty-acid degradation and PUFA metabolism of T-cell subsets in healthy controls, psoriasis and psoriatic arthritis. Front Immunol 2023; 14: 1179877.

39. Xu Y, Wu Y, Xiong Y, Tao J, Pan T, Tan S et al. Ascorbate protects liver from metabolic disorder through inhibition of lipogenesis and suppressor of cytokine signaling 3 (SOCS3). Nutr Metab (Lond) 2020; 17: 17.

40. Xiong X, Liu D, He W, Sheng X, Zhou W, Xie D et al. Identification of gender-related metabolic disturbances in autism spectrum disorders using urinary metabolomics. Int J Biochem Cell Biol 2019; 115: 105594.

41. Ming X, Stein TP, Barnes V, Rhodes N, Guo L. Metabolic perturbance in autism spectrum disorders: a metabolomics study. J Proteome Res 2012; 11(12): 5856–5862.

42. Zhu J, Hua X, Yang T, Guo M, Li Q, Xiao L et al. Alterations in Gut Vitamin and Amino Acid Metabolism are Associated with Symptoms and Neurodevelopment in Children with Autism Spectrum Disorder. J Autism Dev Disord 2022; 52(7): 3116–3128.

43. Liang H, Song K. Elucidating ascorbate and aldarate metabolism pathway characteristics via integration of untargeted metabolomics and transcriptomics of the kidney of high-fat diet-fed obese mice. PLoS One 2024; 19(4): e0300705.

