## Supplementary_figs for "Integrative Analysis of Proteomics and Metabolomics Reveals Impacts of *Sam2* Knockout on Autism Spectrum Disorders"

**Supplementary Fig. 1.**

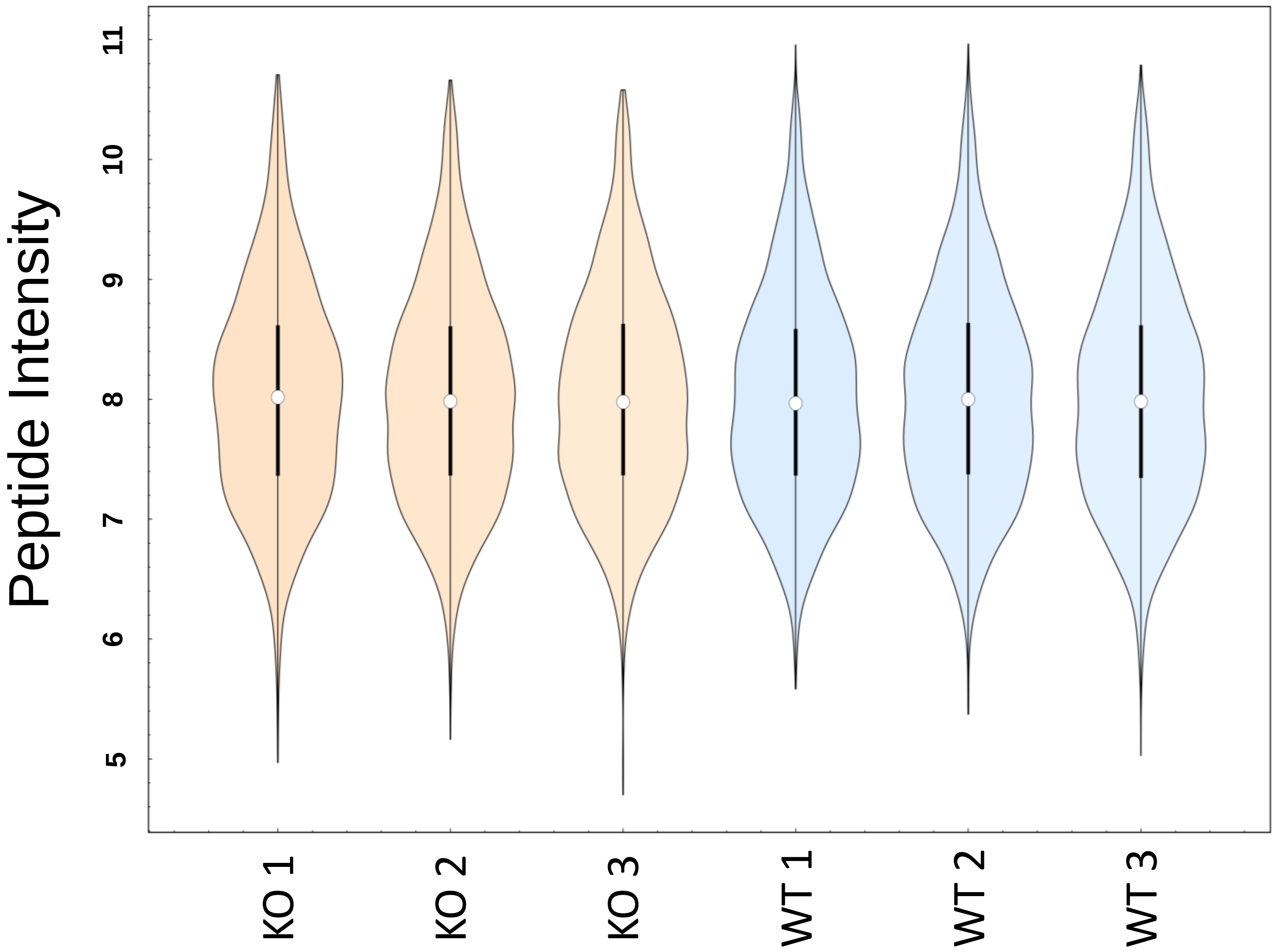

**Supplementary Fig. 2.**

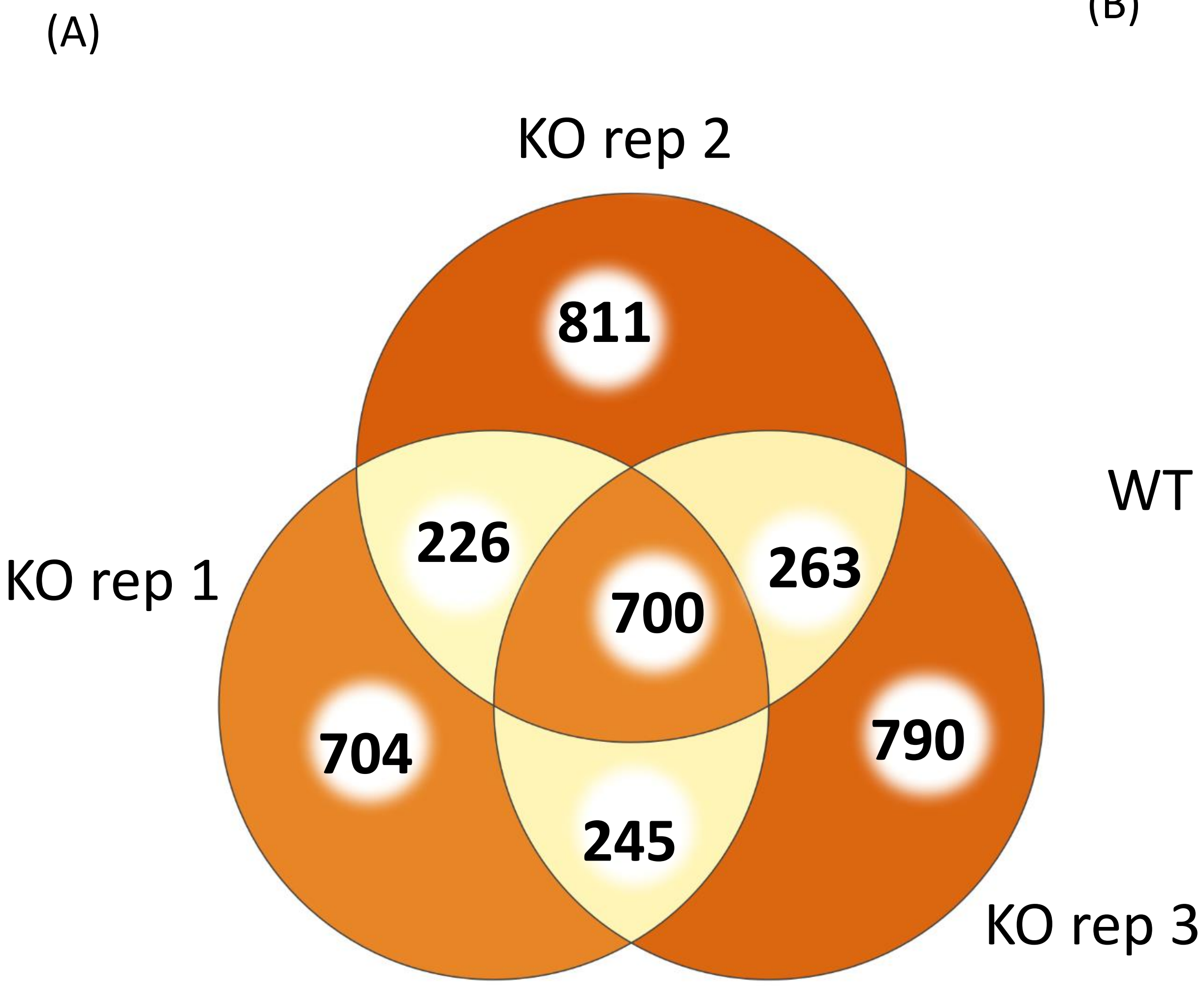


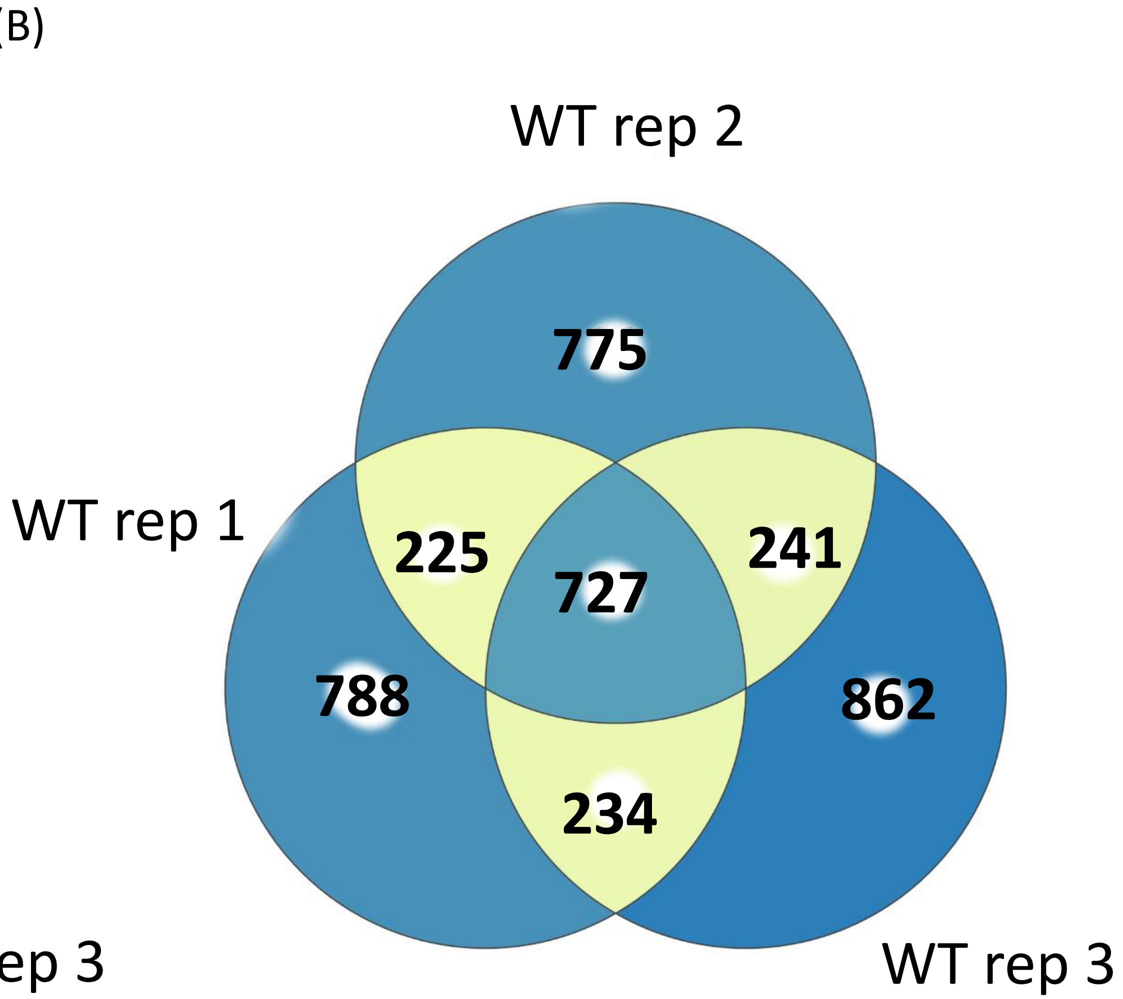

**Supplementary Fig. 3.**

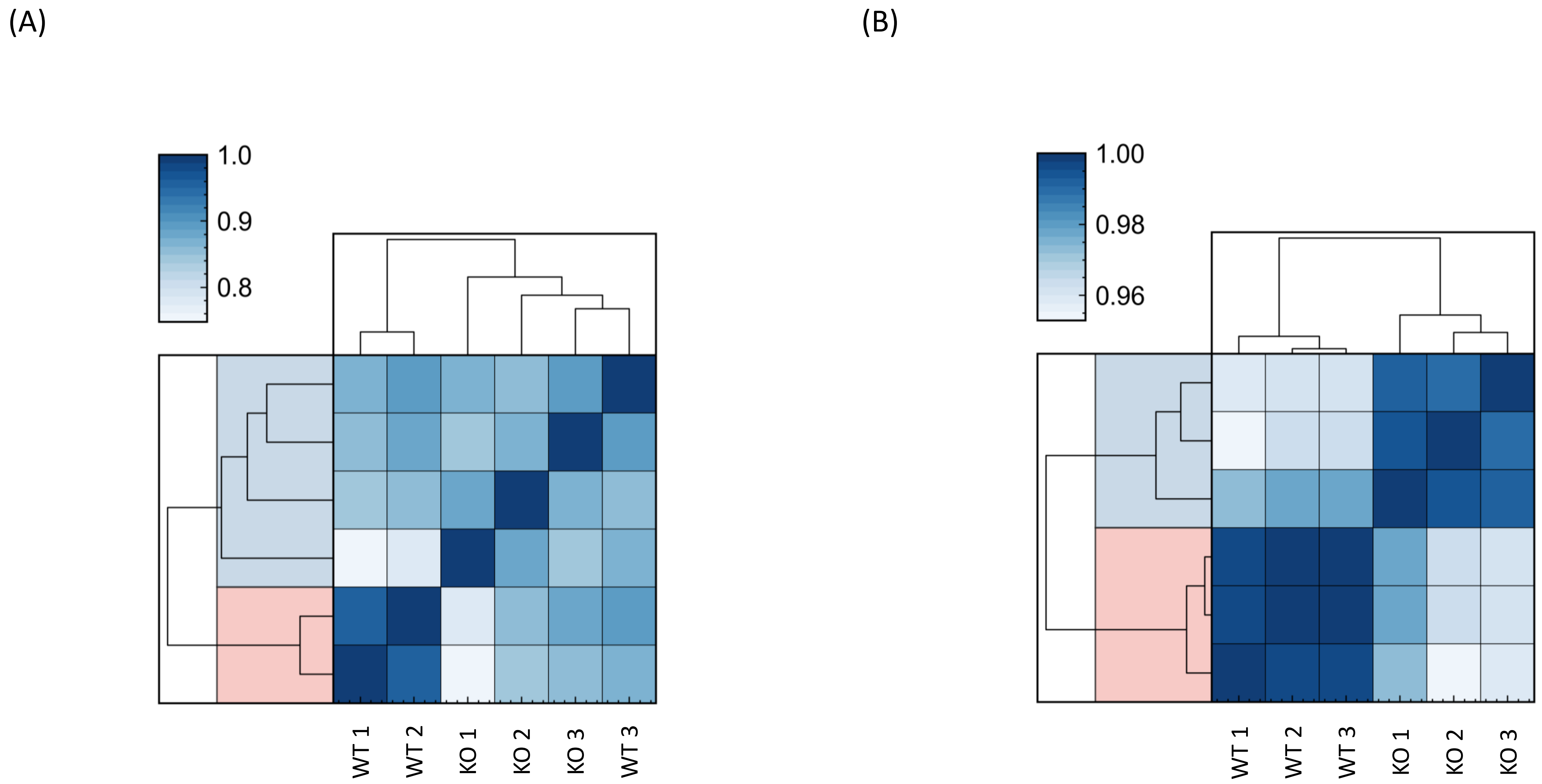

**Supplementary Fig. 3.**

(C) (D)

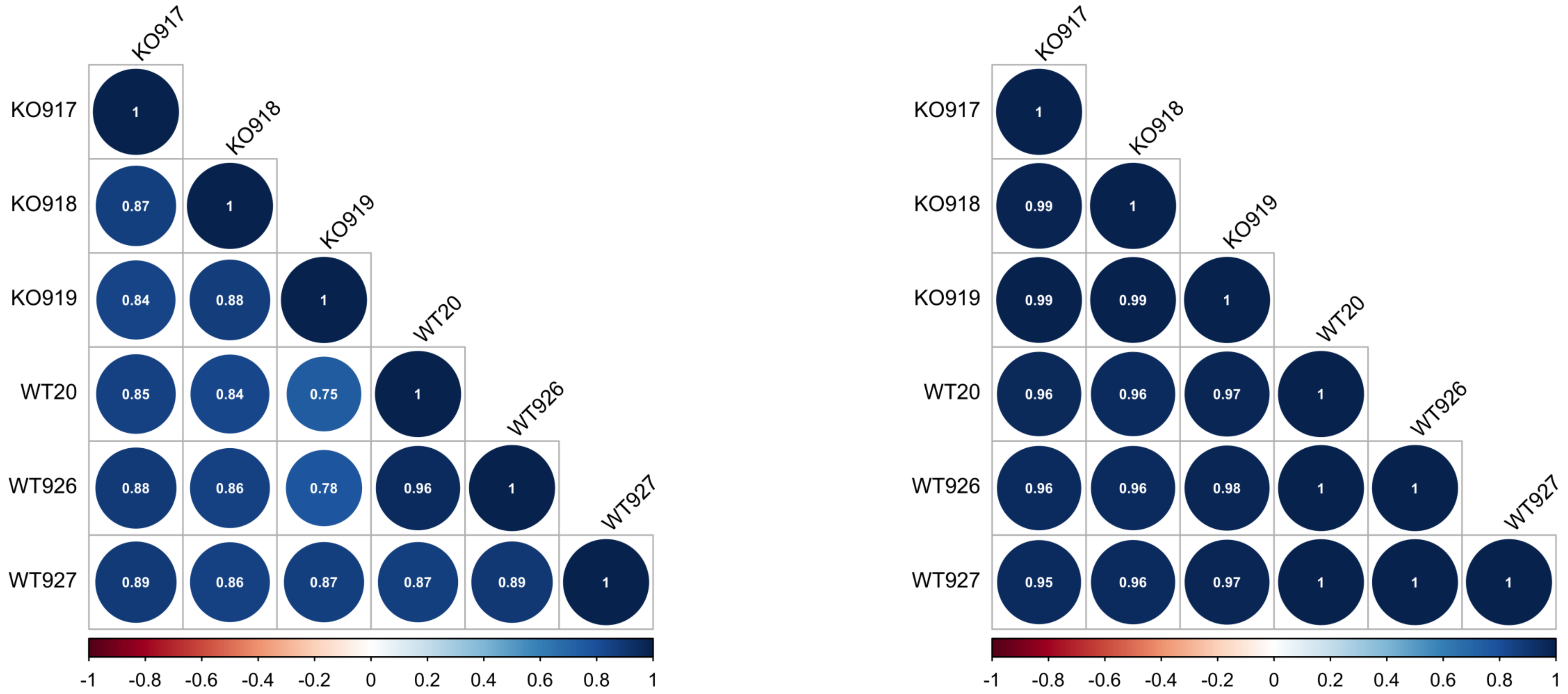

**Supplementary Fig. 4.**

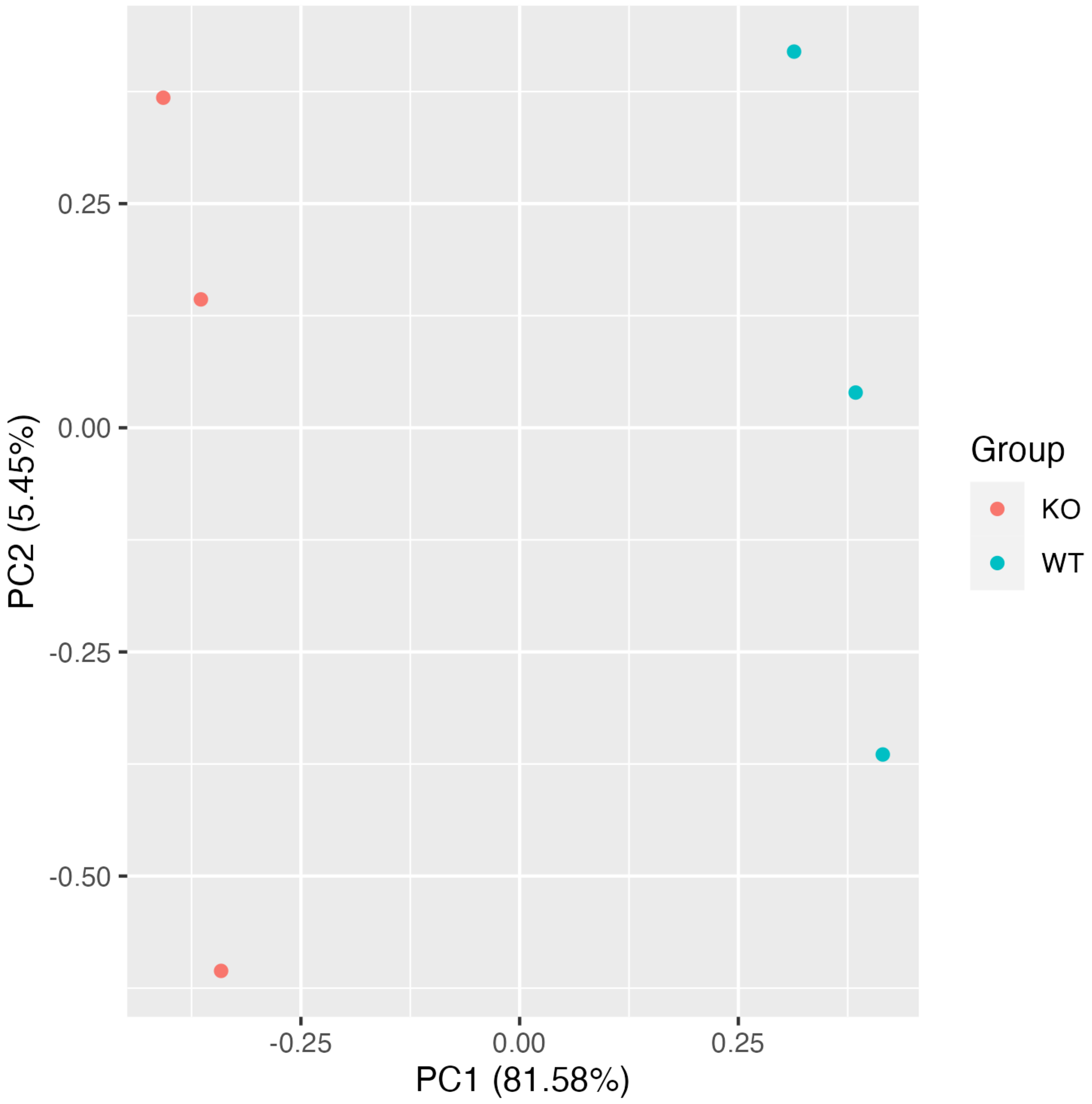

**Supplementary Fig. 5.**

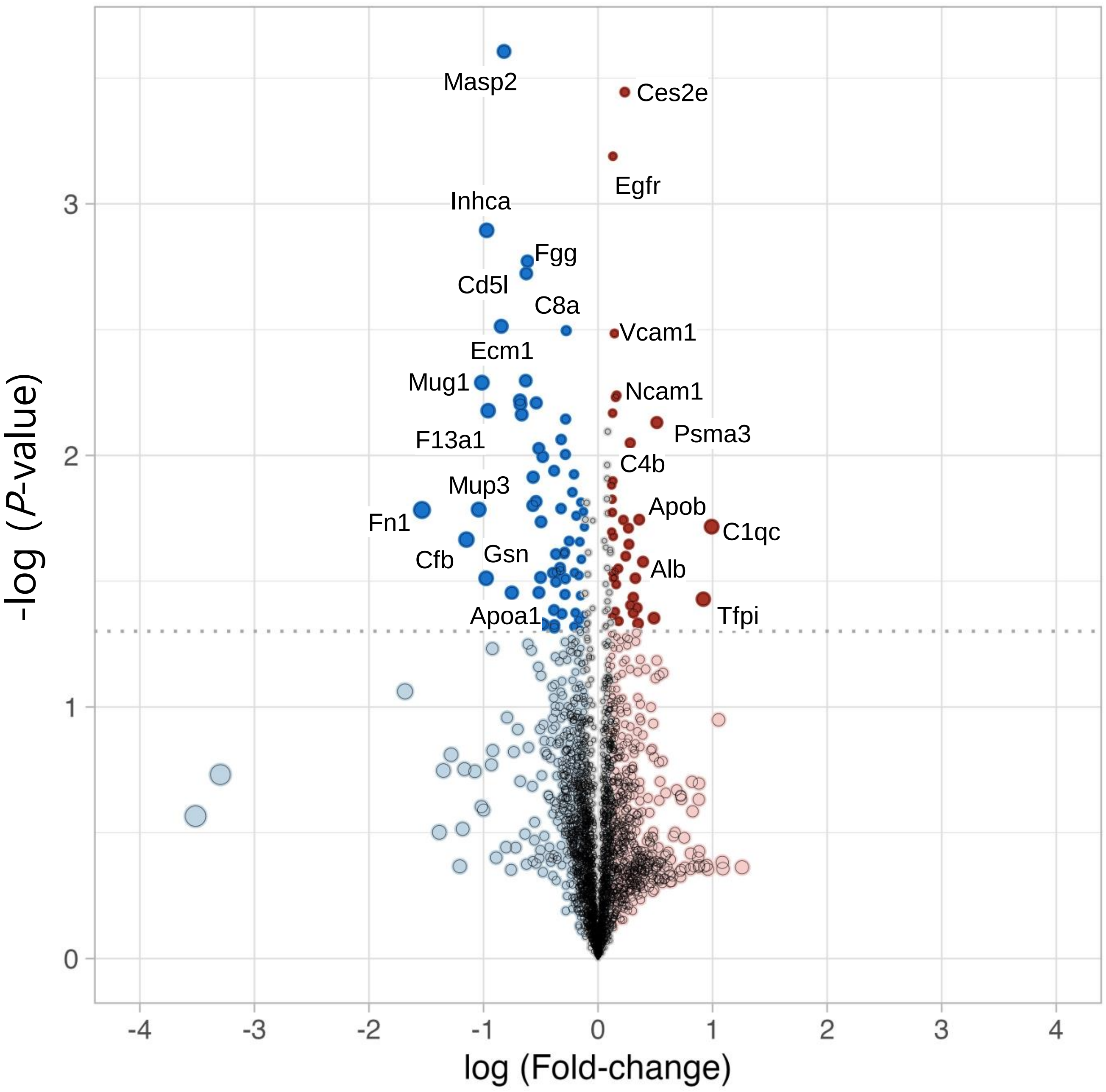

**Supplementary Fig. 6.**

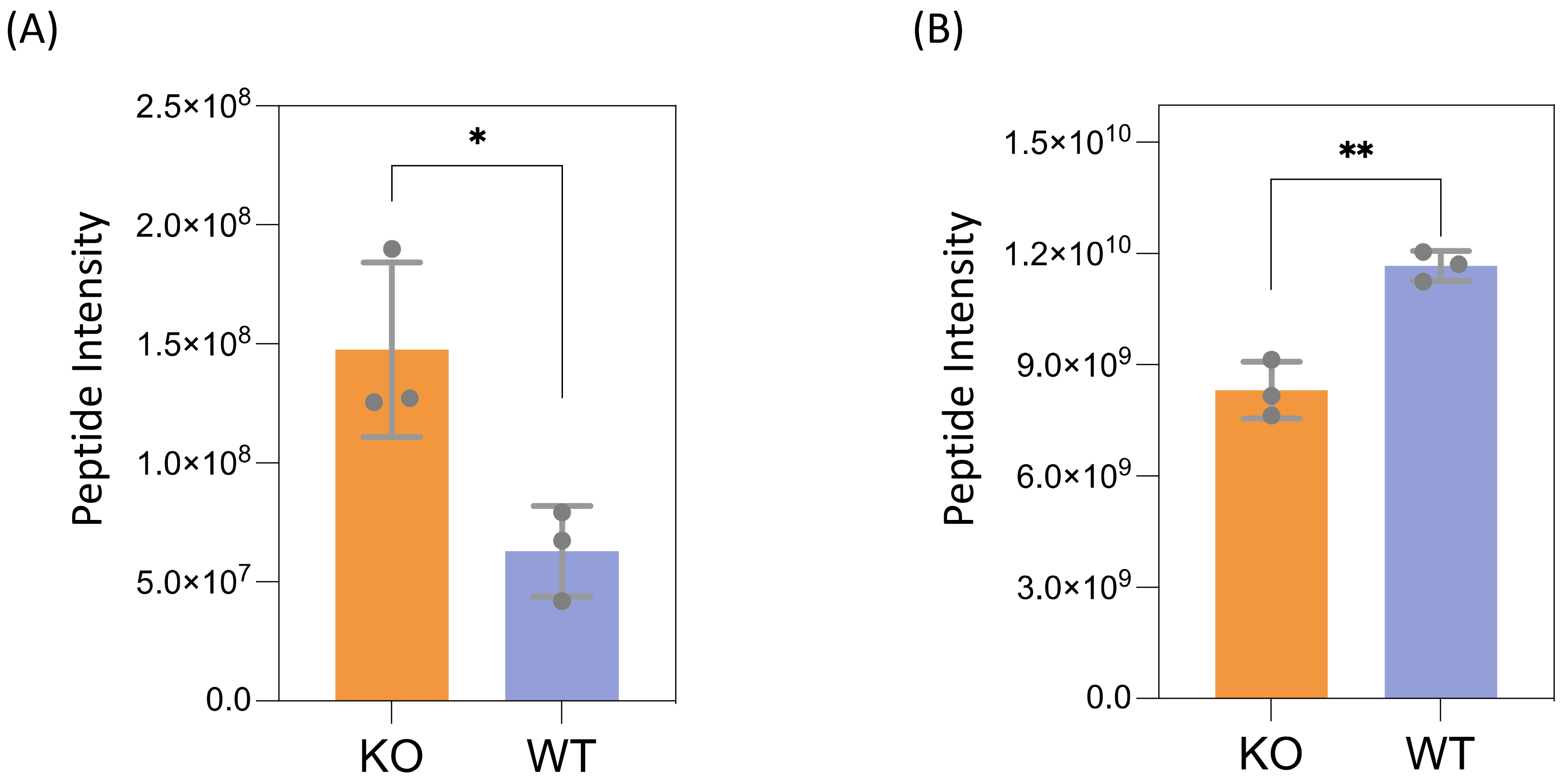


* P-value < 0.05
** P-value < 0.005

**Supplementary Fig. 7.**

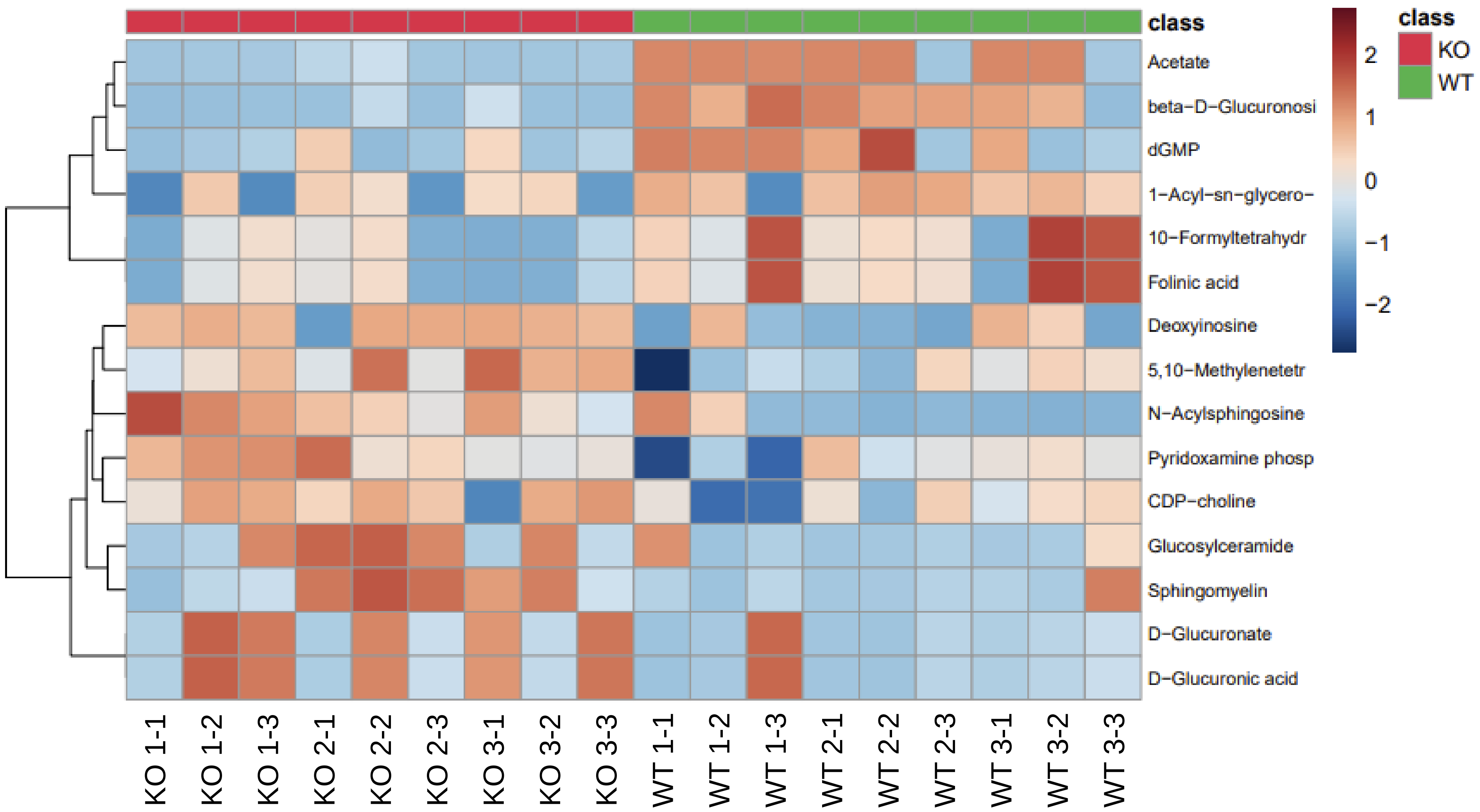

### Supplementary Fig. 8.

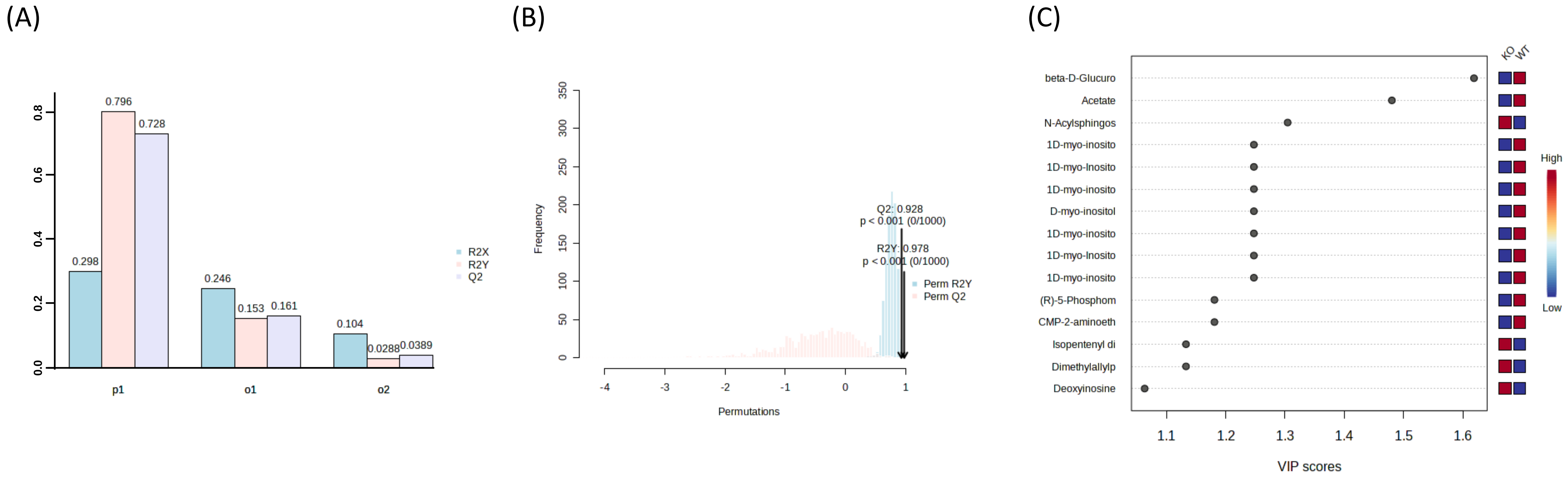
